# An Augmented Aging Process in Brain White Matter in HIV

**DOI:** 10.1101/265199

**Authors:** T. Kuhn, T. Kaufmann, N.T. Doan, L.T. Westlye, J. Jones, R.A. Nunez, S.Y. Bookheimer, E.J. Singer, C.H. Hinkin, A.D. Thames

## Abstract

**Objective:** HIV infection and aging are both associated with neurodegeneration. However, whether the aging process alone or other factors associated with advanced age account for the progression of neurodegeneration in the aging HIV-positive (HIV+) population remains unclear.

**Methods:** HIV+ (n=70) and HIV-negative (HIV-, n=34) participants underwent diffusion tensor imaging (DTI) and metrics of microstructural properties were extracted from regions of interest (ROIs). A support vector regression model was trained on two independent datasets of healthy adults across the adult life-span (n=765, Cam-CAN = 588; UiO = 177) to predict participant age from DTI metrics, and applied to the HIV dataset. Predicted brain age gap (BAG) was computed as the difference between predicted age and chronological age, and statistically compared between HIV groups. Regressions assessed the relationship between BAG and HIV severity/medical comorbidities. Finally, correlation analyses tested for associations between BAG and cognitive performance.

**Results:** BAG was significantly higher in the HIV+ group than the HIV-group *F* (1, 103) = 12.408, p = 0.001). HIV RNA viral load was significantly associated with BAG, particularly in older HIV+ individuals (R^2^ = 0.29, F(7, 70) = 2.66, p = 0.021). Further, BAG was negatively correlated with domain-level cognitive function (learning: r = −0.26, p = 0.008; memory: r = −0.21, p = 0.034).

**Conclusions:** HIV infection is associated with augmented white matter aging, and greater brain aging is associated with worse cognitive performance in multiple domains.

## INTRODUCTION

Advanced age is a major risk factor for cognitive decline and neurodegeneration, including deterioration of white matter (WM) throughout the brain^1, 2, 3^. Older adults with HIV infection are at increased risk for cognitive decline^4, 5^ and WM neurodegeneration^6, 7, 8^ Importantly, the effects of age can be augmented by factors associated with severity of HIV infection and medical comorbidities which may influence the trajectory of HIV infection and cognitive and brain aging.

Neuroimaging-based estimates of the deviance between an individual’s chronological age and predicted brain age - termed brain age gap (BAG) - are sensitive to augmented and/or accelerated brain aging ^9, 10^ While no current study has investigated BAG in WM microstructure using diffusion tensor imaging (DTI), recently, using brain volume, Cole and colleagues^9^ demonstrated that BAG in the volume of brain grey and white matter is greater in HIV-infected individuals than in HIV-controls. Although no significant relationships between BAG and nadir CD4+ or HIV disease duration were reported, the authors investigated white matter volume, rather than white matter microstructure (using DTI) and in a sample of HIV+ participants with undetectable viral loads. Thus, it is unclear to which degree findings of “attenuated brain aging” in HIV^9, 11, 12^ may be mediated by primary or secondary processes related to the infection. It is also unclear what effects these processes may have on WM microstructure. Therefore, this study sought to expand on previous findings by investigating the predicted-BAG in WM microstructure using DTI metrics of white matter patency. This project also sought to further elucidate the effect of HIV severity indices and non-HIV-related medical comorbidities on accentuated WM aging.

Here, we used a machine learning approach to quantify brain aging based on DTI. A support vector machine model trained in a large and independent training-set of healthy controls was used to predict age in HIV+ and comparable HIV-individuals. We tested for group differences in BAG between HIV+ and HIV-, and for associations with cognitive function and HIV disease factors as well as medical comorbidities within the HIV+ group.

## METHODS

### PARTICIPANTS

Participants included a testing dataset of 104 (72 HIV+ (confirmed by serologic testing); 32 HIV-) adults (M_age_= 50.17; SD = 12.82) who were enrolled as part of a larger study (K23 MH095661; PI: A.D.T.). All participants who had DTI imaging data available were included in this study. All procedures were in accordance with the Declaration of Helsinki, reviewed and approved by the University of California, Los Angeles (UCLA) Institutional Review Board prior to enrollment and all participants provided written informed consent.

The training set used for the age prediction model comprised 765 healthy community dwelling individuals aged 20-78 years sampled from two different cohorts, including the ongoing STROKEMRI study at the University of Oslo (UiO; n=177, M_age_= 57.59; SD = 15.05, 60 % female, PI: L.W.) and the Cambridge Centre for Ageing and Neuroscience ^13^ (Cam-CAN) (n=588, M_age_= 52.31; SD = 17.38, 50 % female) from the Cambridge Center for Ageing and Neuroscience ^14^ (CamCAN). Data used in the preparation of this work were obtained from the CamCAN repository (available at http://www.mrc-cbu.cam.ac.uk/datasets/camcan/)^13, 14^. All procedures were in accordance with relevant IRBs.

#### PSYCHIATRIC ASSESSMENT

In the test sample, the Structured Clinical Interview (SCID) for DSM-IV ^15^, and structured questionnaires were used to screen for neurological, psychiatric, and medical confounds including: history of seizure disorder or other neurologic disorder; history of concussion or traumatic brain injury sufficient to warrant medical attention; history of Axis I psychiatric disorder or current substance use disorder (SCID-IV diagnostic criteria); current prescriptions for psychotropic medication, except for anxiolytics and antidepressants; current substance dependence or stimulant use, comorbid infection (e.g. Hepatitis C), HIV-associated CNS opportunistic infection (e.g. CNS toxoplasmosis) or CNS neoplasm. Participants were also screened for contraindications to MRI.

#### COGNITIVE ASSESSMENTS

In the test sample, participants completed a comprehensive neuropsychological test battery used in prior studies^16^, which assessed neurocognitive function at both the global and domain level. Six cognitive domains were measured: (1) <u>Processing Speed</u> -Wechsler Adult Intelligence Scale—Fourth Edition (WAIS-IV) Digit Symbol and Symbol Search subtests^17^, Trail Making Test—Part A^18^, and Stroop—Color Naming and Word Reading^19^; (2) <u>Learning</u> - Hopkins Verbal Learning Test—Revised^20^ and Brief Visuospatial Memory Test—Revised^21^; (3) <u>Memory</u> - Hopkins Verbal Learning Test—Revised^20^ and Brief Visuospatial Memory Test—Revised^20, 21^ (delayed recall); (4) <u>Language/Verbal Fluency</u> - Controlled Oral Word Association Test^22^ (FAS and Animals); (5) <u>Executive Function</u> - WAIS-IV Letter-Kuhn, T., Ph.D. Number Sequencing subtest^17^, Trail Making Test—Part B^18^, and Stroop-Color-Word Interference Test^19^; and (6) <u>Motor Speed</u> - Grooved Pegboard test^23^ (dominant and non-dominant hands).

We converted raw test scores into within-sample z scores and then averaged them to create neurocognitive domain z scores. We calculated the global neurocognition score by averaging the z scores from all of the neuropsychological test variables. Given that the relationship between age and neurocognitive performance in HIV is a primary aim of this study, within-sample z scores were computed instead of demographically-adjusted T scores.

#### IMMUNE STATUS ASSESSMENT

In the testing dataset, HIV+ participants self-reported nadir CD4+ and lifetime highest viral load were used to assess past immune status. Participants also underwent venipuncture to test current CD4+ and HIV viral load. HIV duration was calculated as the number of years since the participant’s self-reported HIV diagnosis. Next, participants were classified as either ‘pre-HAART’ (highly active antiretroviral therapy) or ‘post-HAART’ based on whether their initial HIV diagnosis was before or after 1996^24^. Further, a “medical comorbidity burden’ index score was computed from the medical history taken during the routine interview all participants completed during data collection. Participants were assigned a ‘1’ if they endorsed a history of each of the following medical conditions: cerebrovascular risk factors including hypertension, heart failure, COPD, anemia, diabetes; endocrine dysfunction including thyroid disease, testosterone therapy, estrogen therapy; kidney disease. Participants were assigned a ‘0’ for all medical conditions they did not endorse. The ‘medical comorbidity burden’ score was then computed as a sum of these conditions, resulting in a scale ranging from 0 (no medical comorbidities) to 9 (endorsed all medical comorbidities identified).

### MRI ACQUISITION AND ANALYSIS

Diffusion weighted imaging (DWI) scans were collected from HIV+ individuals and demographically-matched HIV-seronegative controls at the University of California, Los Angeles, all of whom comprised the testing dataset. This MR data was collected using a 3T Siemens Trio scanner (Siemens, Germany) at the UCLA Center for Cognitive Neuroscience (CCN). 64 diffusion-weighted volumes (*b*=1,000 s/mm^2^) and 6 non-diffusion-weighted volumes were obtained using a single shot spin-echo echo planar imaging (EPI) sequence with 60 × 2.0 mm axial slices (no gap), flip angle = 90, TR = 9000 ms, TE = 93 ms, voxel size = 2.0 × 2.0 × 2.0 mm, b-shells = 0, 1000, scan time 576 s.

Training data including MRI data from UiO and Cam-CAN. UiO MRI data were collected on a General Electric 750 3T scanner (General Electric, United States) at Oslo University Hospital. 60 diffusion-weighted volumes (*b*=1,000 s/mm^2^) and 5 non-diffusion-weighted volumes were obtained using a single shot spin-echo EPI sequence with 67 × 2.0 mm axial slices (no gap), flip angle = 90, TR = 8150 ms, TE = 83.1 ms, voxel size = 2.0 × 2.0 × 2.0 mm, b-shells = 0, 1000, 60 directions, scan time 530 s. The Cam-CAN DWI images were collected on a Siemens TIM Trio 3T scanner (Siemens, Germany) at the Medical Research Counsel (UK) Cognition and Brain Sciences Unit (MRC-CBSU). 60 diffusion-weighted volumes (30 with *b*=1,000 s/mm^2^ and 30 with *b*=2,000 s/mm^2^) and 3 non-diffusion-weighted volumes were obtained using a twice refocused spin-EPI with 66 × 2.0mm axial slices (no gap), TR = 9100 ms, TE = 104 ms, voxel size = 2.0 × 2.0 × 2.0 mm, scan time 573 s. Only the b=0 and b=1000 shells were used for DTI analysis in the present study. All DWI images were quality controlled and visually inspected prior to being preprocessed and analyzed.

DWI scans from UCLA, UiO and Cam-CAN were processed simultaneously through the same pipeline to harmonize imaging methods across sites. All imaging data were processed using FMRIB software Library^25^ (FSL, www.fmrib.ox.ac.uk.fsl). DWI data was motion and eddy current corrected using EDDY^26^, skull stripped using BET^27^, and then diffusion tensors were fit to the data using dtifit in FSL. Tract-Based Spatial Statistics^28^ (TBSS) was used to generate a WM skeleton comprised of WM voxels shared by all participants. This WM skeleton was applied to each participant’s individual DTI maps and mean Fractional anisotropy (FA), axial (AD), radial (L1) and mean diffusivity (MD) were extracted from various regions-of–interest (ROIs) based on the intersection between the TBSS skeleton and labels defined in probabilistic anatomical atlases^29, 30, 31^ in addition to a global average across the skeleton. The full list of ROIs from which these DTI metrics were extracted follows: anterior, posterior and superior corona radiata; anterior, posterior and retrolenticular portions of the internal capsule; anterior and posterior thalamic radiation; sagittal stratum; external capsule; WM underlying the cingulate bundle; WM underling the hippocampus; inferior, middle and superior cerebellar peduncles; cerebral peduncle; pontine; fornix; stria terminalis; corticospinal tract; medial lemniscus; inferior and superior longitudinal fasciculus; superior fronto-occipital fasciculus; uncinate fasciculus; genu, body, tapetum and splenium of the corpus callosum.

### STATISTICAL ANALYSIS

#### GROUP DEMOGRAPHICS

Participant characteristics (e.g. age, education, past drug use) between HIV+ and HIV-groups were compared using one-way analysis of variance (ANOVA). Group differences in dichotomous factors (e.g. sex, ethnicity, urinalysis results) were assessed using chi-square analyses. We used p< .05 as our cutoff for statistical significance for these demographic analyses.

#### AGE PREDICTION AND BRAIN AGE GAP

UiO and Cam-CAN data were used to train a support vector regression model (SVR) to predict participant age using FA, L1, RD and MD from atlas-derived ROIs (the exact same regions described above^29, 30, 31^) as features. Similar methods have been employed using imaging data previously^9, 10, 32^, including using DTI to assess participant age in a healthy cohort^33^. SVR was conducted in Matlab (https://mathworks.com/help/stats/fitrsvm.html) using the implementation “fitrsvm” with a linear kernel, automatic hyperparameter tuning and Sequential Minimal Optimization. Given that multiple MR scanners were used to collect the HIV and training data, scanner was used as a regressor on the features to control for interscanner variability. The model accuracy was validated using 10-fold cross-validation on the training set. After successful validation, the trained SVR was used to predict age of participants in the independent UCLA sample (HIV+ / HIV-). For each individual, BAG was computed by subtracting the participant’s predicted brain age by their chronologic age. Univariate analysis of covariance (ANCOVA) then compared BAG between UCLA HIV serostatus groups, controlling for chronological age and sex.

#### ASSOCIATIONS BETWEEN BRAIN AGE GAP IMMUNE STATUS AND COGNITIVE PERFORMANCE

Within HIV+ subjects,_a stepwise hierarchical linear regression was conducted to investigate the relationship between BAG and chronological age, nadir CD4+, lifetime highest HIV RNA viral load and medical comorbidity index, and all possible interactive relationships between age and other dependent variables using a stepwise entry model.

Correlation analyses were used to investigate the relationship between BAG and cognitive performance in the UCLA HIV+ group. Correlations were conducted between BAG and individual cognitive domains (e.g. attention, memory) as well as global cognitive performance. We report both bivariate correlations as partial correlations covarying for premorbid intellectual ability (Wide Range Achievement Test, 4^th^ Edition (WRAT-4)^34^). False discovery rate (FDR) was used to correct for multiple comparisons.

## RESULTS

### DEMOGRAPHIC GROUP COMPARISON

In the test dataset, the HIV+ and HIV-seronegative groups did not significantly differ on age, years of education, ethnicity, or sex (Table 1). The HIV+ group had significantly higher medical comorbidity burden (i.e. greater number of medical comorbidities) than the HIV-control group (X^2^ = 7.39, p = 0.007). None of the participants tested positive for barbiturates, cocaine, methamphetamine, phencyclidine, or MDMA. Significantly more HIV+ participants tested positive for prescribed benzodiazepines (X^2^ = 5.93, p = 0.015) than did HIV-participants. The HIV serostatus groups did not differ on current alcohol or substance abuse, past (i.e. self-reported lifetime) substance dependence, past substance abuse, past alcohol dependence or past alcohol abuse (all p’s > 0.10). Participants were not included in the study if they reported previous methamphetamine abuse or dependence.

Within the HIV+ group, participants diagnosed in the pre-HAART era were significantly older (M = 56.85, SD = 8.93) than those diagnosed in the post-HAART era (M = 48.24,SD = 12.16) (F(1, 70) = 8.22, p = 0.006). The pre-and post-HAART HIV+ groups did not significantly differ on current CD4+, current HIV viral load, nadir CD4+ or lifetime highest viral load (all p’s >0.1). Further, there was not a significant difference between the younger (50 years old and below) and older (51 years old and above) HIV+ participants on current CD4+, current HIV viral load, nadir CD4+ or lifetime highest viral load (all p’s >0.1). Table 1 and Figure 1 provide additional detail on group demographics.

**Table 1.**
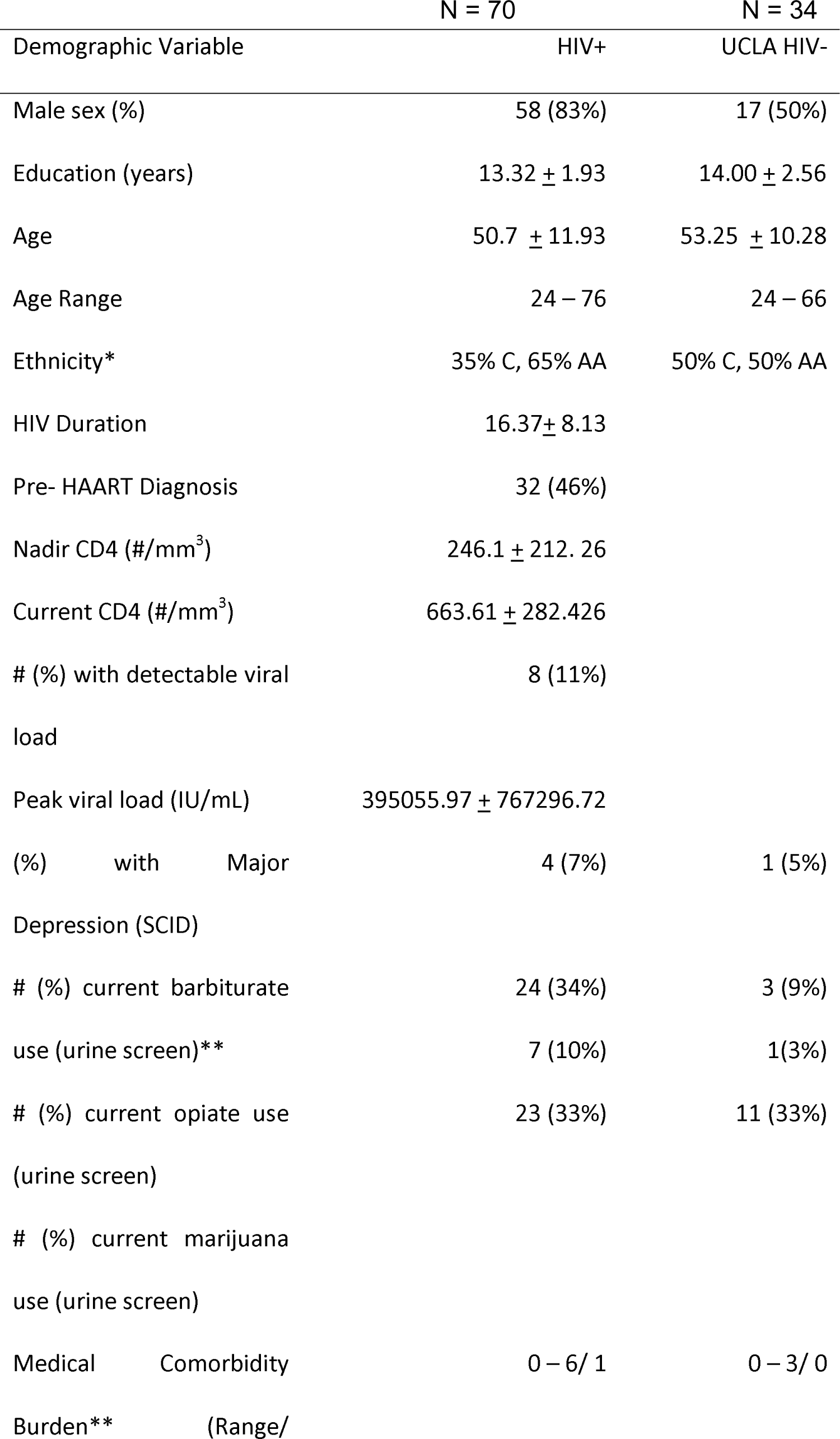
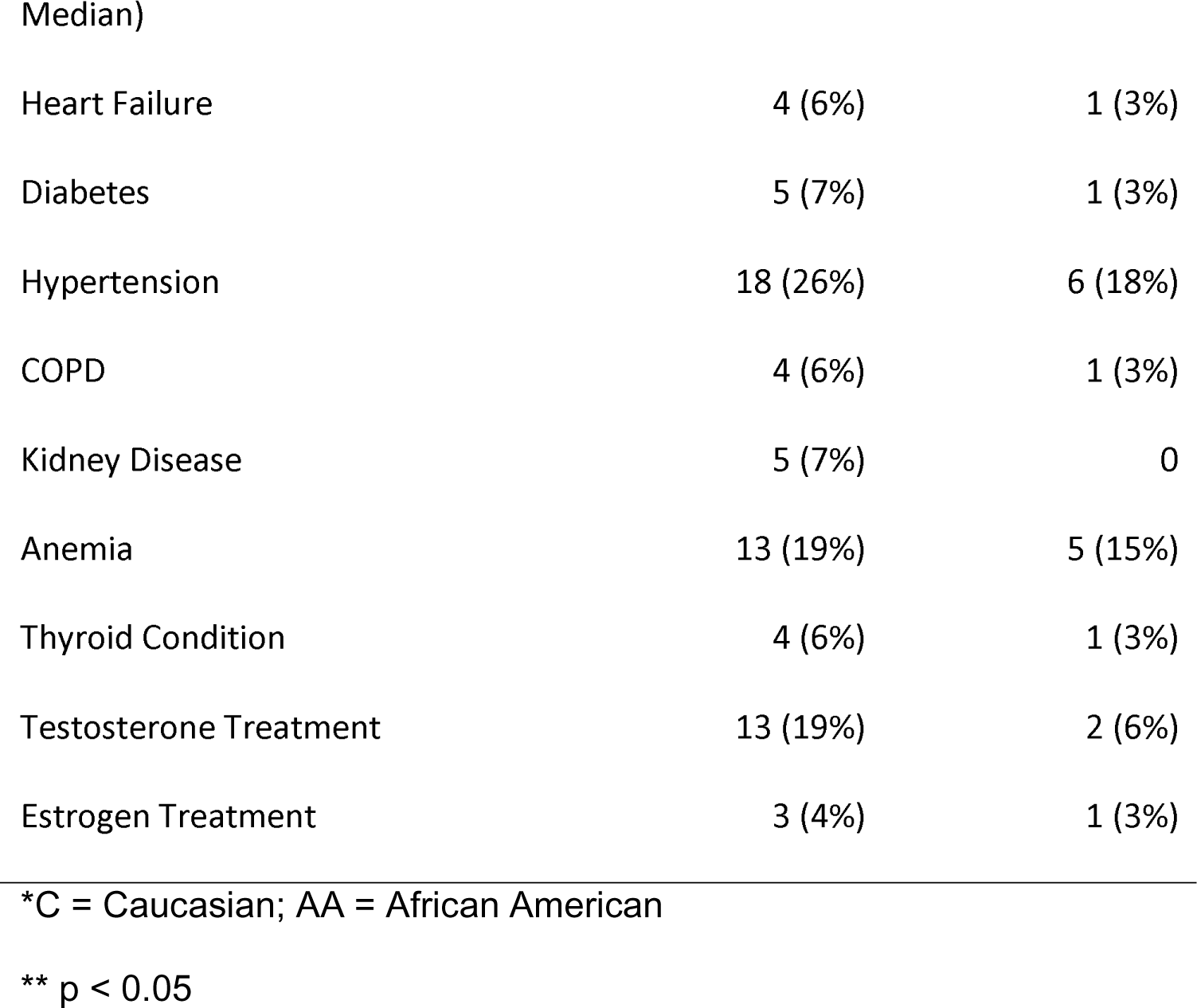
DEMOGRAPHIC COMPARISON BETWEEN HIV+ & HIV-GROUPS.

| Demographic Variable | N = 70 | N = 34 |
| --- | --- | --- |
|  | HIV+ | UCLA HIV- |
| Male sex (%) | 58 (83%) | 17 (50%) |
| Education (years) | 13.32 ± 1.93 | 14.00 ± 2.56 |
| Age | 50.7 ± 11.93 | 53.25 ± 10.28 |
| Age Range | 24 – 76 | 24 – 66 |
| Ethnicity* | 35% C, 65% AA | 50% C, 50% AA |
| HIV Duration | 16.37 ± 8.13 | |
| Pre- HAART Diagnosis | 32 (46%) |  |
| Nadir CD4 (#/mm<sup>3</sup>) | 246.1 ± 212. 26 | |
| Current CD4 (#/mm<sup>3</sup>) | 663.61 ± 282.426 | |
| # (%) with detectable viral load | 8 (11%) |  |
| Peak viral load (IU/mL) | 395055.97 ± 767296.72 | |
| (%) with Major Depression (SCID) | 4 (7%) | 1 (5%) |
| # (%) current barbiturate use (urine screen)** | 24 (34%) | 3 (9%) |
| # (%) current opiate use (urine screen) | 7 (10%) | 1(3%) |
| # (%) current marijuana use (urine screen) | 23 (33%) | 11 (33%) |
| Medical Comorbidity | 0 – 6/ 1 | 0 – 3/ 0 |
| Burden** (Range/ |  |  |
| Median) |  |  |
| Heart Failure | 4 (6%) | 1 (3%) |
| Diabetes | 5 (7%) | 1 (3%) |
| Hypertension | 18 (26%) | 6 (18%) |
| COPD | 4 (6%) | 1 (3%) |
| Kidney Disease | 5 (7%) | 0 |
| Anemia | 13 (19%) | 5 (15%) |
| Thyroid Condition | 4 (6%) | 1 (3%) |
| Testosterone Treatment | 13 (19%) | 2 (6%) |
| Estrogen Treatment | 3 (4%) | 1 (3%) |

\*C = Caucasian; AA = African American
\*\* p < 0.05

**Figure 1.**
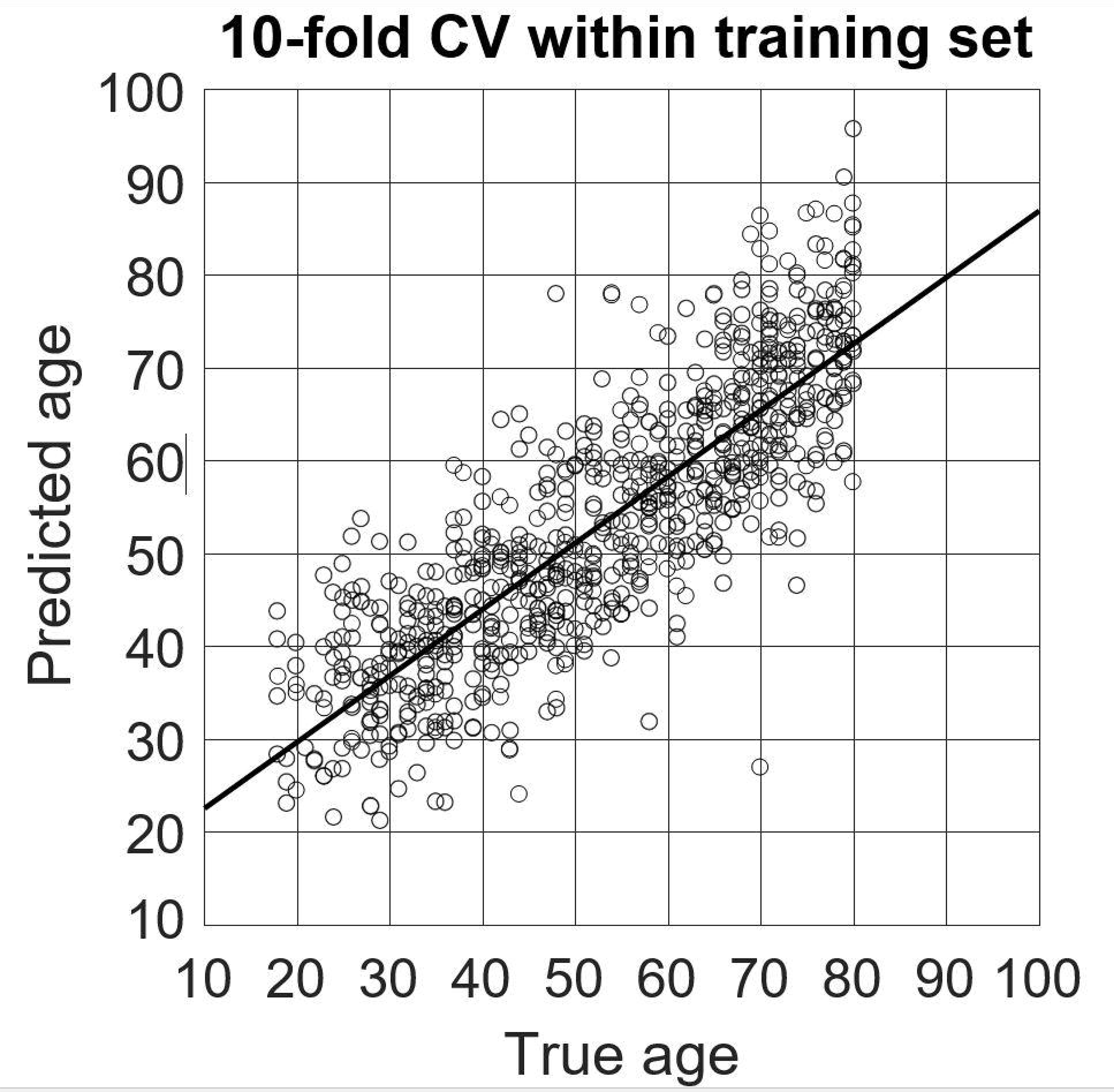
AGE OF GROUPS

### AGE PREDICTION

Predicted brain age, derived from DTI metrics extracted from ROIs (previously described), was strongly correlated with chronological age in the training set (r = 0.84, R^2^ = 0.70, MAE = 7.39, RMSE = 10.64, p <.0001, 10-fold cross-validation), indicating successful tuning of the trained SVR model. Applied to UCLA testing data, the model successfully predicted brain age in the independent sample, both in the UCLA HIV- (r = 0.78, R^2^ = 0.61, MAE = 7.64, RMSE = 9.43, p < 0.0001) and HIV+ groups (r = 0.64, R^2^ = 0.41, MAE = 9.48, RMSE = 12.004, p < 0.0001). Participant true chronological age was not correlated with prediction error (r = -.10, p = 0.31). Figure 2 depicts age prediction results. Figure 3 is a scatterplot depicting the 10-fold cross-validation of the SVR within the training data set.

**Figure 2.**
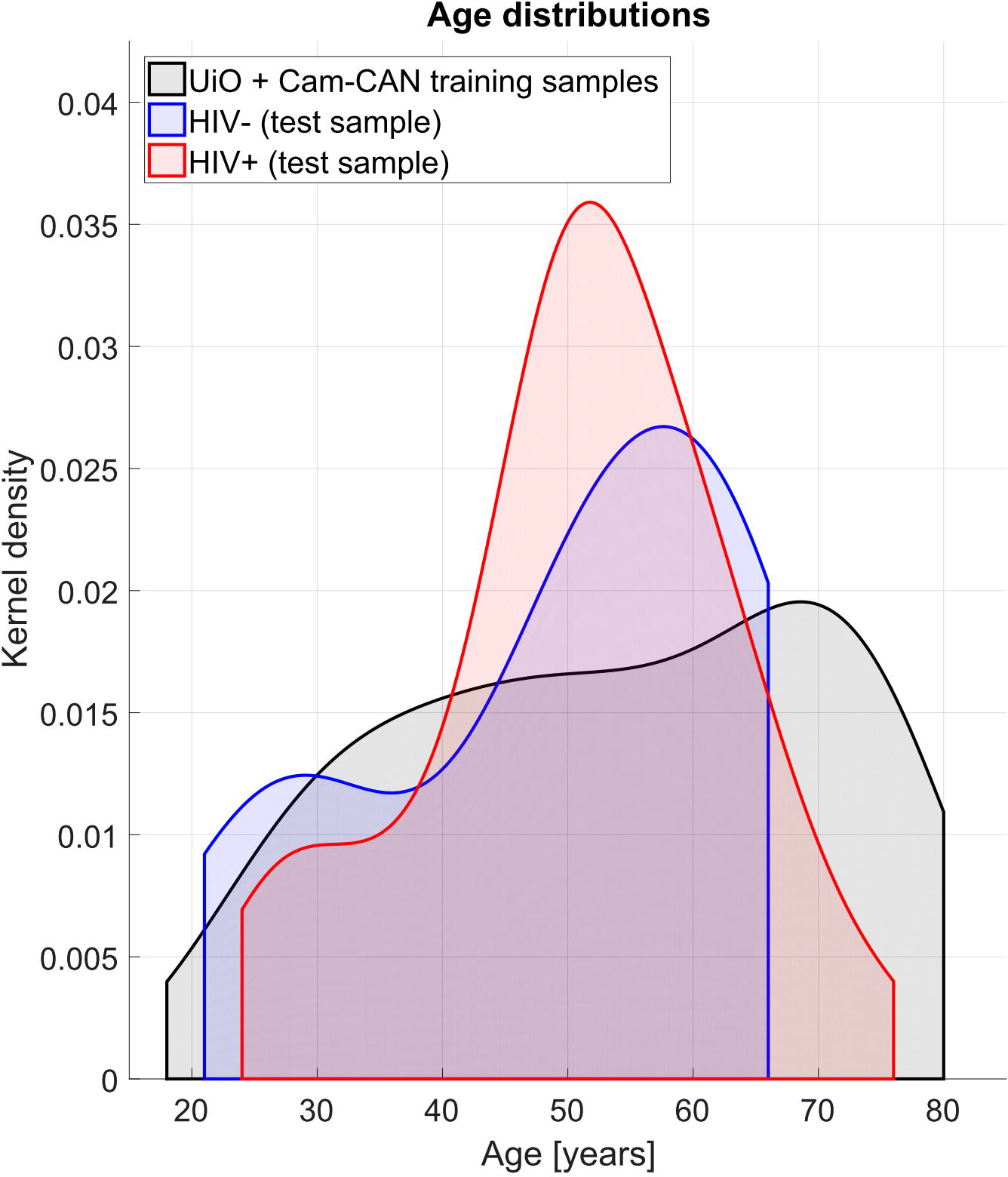
SVR-BASED AGE PREDICTION

**Figure 3.**
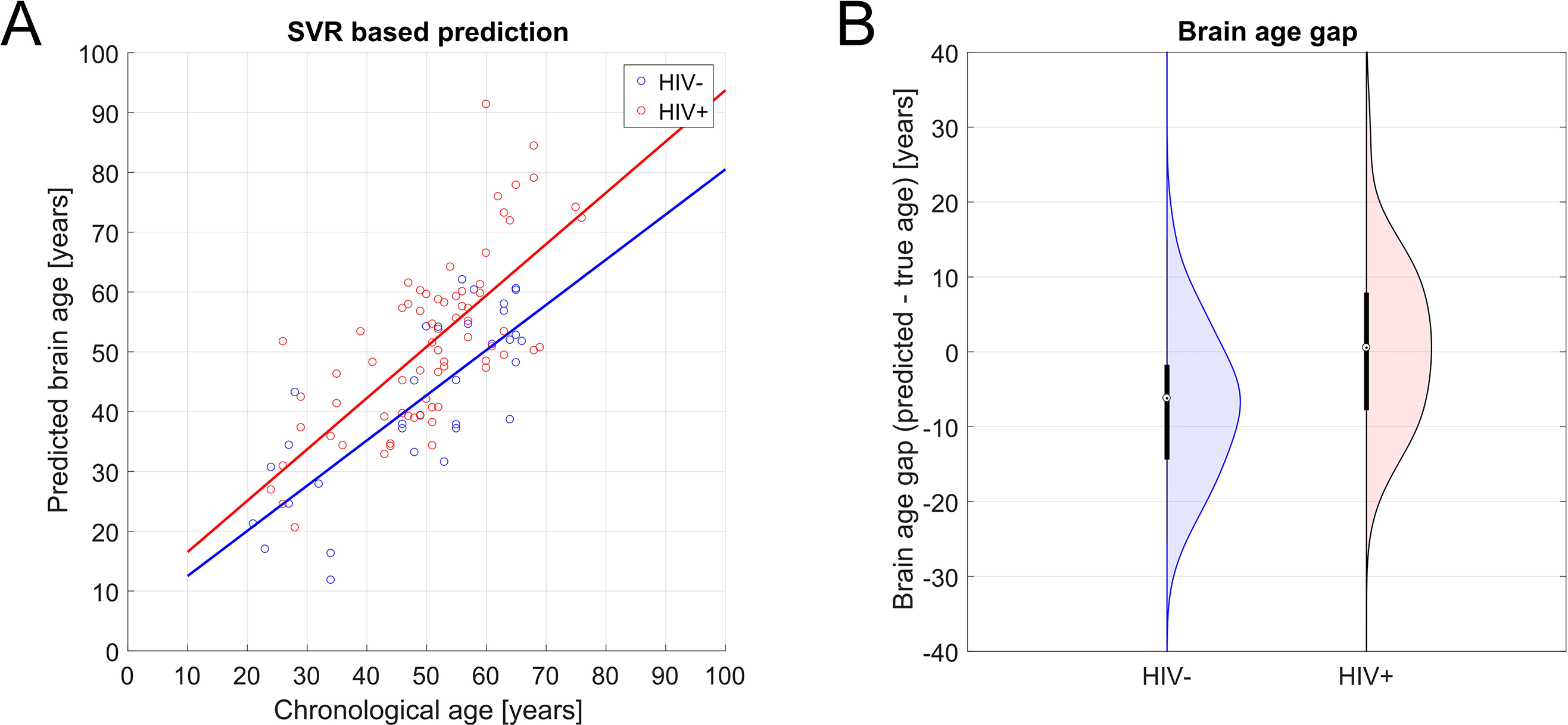
TEN-FOLD CROSS-VALIDATION WITHIN TRAINING SET

In the testing dataset, HIV+ individuals showed higher differences between their brain age and their chronological age than HIV-counterparts *(F* (1, 103) = 12.408, p = 0.001, partial η^2^ = 0.21). As participant chronologic age increased, BAG also increased (F (39, 103) = 5.57, p = 0.010, partial η^2^ = 0.37). There was no age*HIV interaction on BAG (p > 0.05).

### NEUROCOGNITION, IMMUNE STATUS & BRAIN AGE GAP

All results reported below withstood FDR correction.

A regression that used chronological age, HIV duration, pre-versus post-HAART diagnosis, medical comorbidity burden, nadir CD4+ count and lifetime highest HIV RNA viral load (log transformed) as predictors of BAG (R^2^ = 0.38, F(6, 70) = 4.13, p = 0.004) revealed associations between BAG and chronological age (β = −0.38, p = 0.006) and highest HIV RNA viral load (β = 0.34, p = 0.004) in the HIV+ group. No other dependent variable was significantly associated with BAG (all p’s > 0.10).

Next, the following step in the stepwise hierarchical regression resulted in the inclusion of the significant interactive effect of chronological age and lifetime highest HIV RNA viral load. No other interactive effects were significant (all p’s > 0.10). There was a significant interactive effect of chronological age and highest HIV RNA viral load (β = 0.23, p = 0.033) on BAG (R^2^ = 0.29, F(7, 70) = 2.66, p = 0.021). Both main effects for chronological age (β = −0.42, p = 0.007) and highest HIV RNA viral load (β = 0.22, p = 0.035) remained significant. In other words, older HIV+ participants who reported a history of higher viral load, had the greatest discrepancy between their estimated brain age and chronological age.

We also found significant associations between BAG and learning (r = −0.26, p = 0.008) and between BAG and memory (r = −0.21, p = 0.034). When controlling for premorbid intellectual ability (WRAT), both associations remained (learning: r = −0.27, p = 0.008; memory: r = −0.20, p = 0.041).

## DISCUSSION

This study used machine learning along with a large training data set of normal WM aging to examine HIV-associated WM microstructural alterations and related this WM degeneration to cognitive impairment. Using an SVR trained with this healthy aging cohort to reliably predict participant age based on metrics of WM microstructure, we found that the brain WM age difference (BAG) was significantly higher in our HIV+ group than in the highly comparable HIV-group. Further, BAG widened with increasing age, suggesting that advancing age is a risk factor for neurodegeneration. Additionally, larger BAG was associated with worse cognitive performance, indicating that this neurodegeneration may be related to deleterious changes in cognition. Although chronological age was not significantly correlated with prediction error, confirmatory analyses verified that these significant associations between BAG and cognitive performance remained after controlling for participant true age.

Importantly, in our sample in which 11% of the HIV+ participants evidenced a detectable HIV RNA viral load, current CD4 and detectable viral load were not related to BAG. Conversely, highest lifetime HIV RNA viral load, which was not the current viral load for any participant, was related to BAG in the HIV+ group, even after controlling for factors such as HIV duration, pre-versus post-HAART diagnosis and medical comorbidities. Highest lifetime viral load also was not different between the younger and older HIV+ groups. Furthermore, older HIV+ participants who reported high plasma viral load had the greatest discrepancy between their estimated brain age and chronological age, suggesting that history of high viral burden contributes to accentuated brain aging. These findings suggest that the impact of early disease burden, even among a sample comprised of 11% of participants with currently detectable HIV RNA viral load, has adverse effects on brain/cognition as HIV+ individuals age. This provides further support of previous findings^9,^ that demonstrated this augmented aging effect in a sample with no participants with detectable HIV RNA. The relationship of the past plasma viral load to current brain reservoirs of HIV are not known, and we can only speculate that persons with a higher plasma viral burden in blood may have also acquired more viral seeding in brain; this may be the stimulus for greater neuroinflammation and more neurodegeneration over years of exposure. Taken together, these findings may suggest that HIV is associated with an augmented aging process in WM which is itself associated with lower cognitive performance.

The mechanisms by which HIV and age result in augmented neurodegeneration are unclear. Holt and colleagues^6^ suggest two potential explanations regarding the relationship between HIV infection and increased brain aging. First, the increased brain age (BAG) may be explained by premature WM aging resulting from the virus facilitating neurodegenerative processes^35^, such as axonal injury, loss of axonal density, reduced patency of axons. Alternatively, advanced age may increase the effects of the virus on the CNS, thereby creating a synergistic interaction effect between HIV and aging^6^. Our finding that lifetime highest HIV RNA viral load, particularly in the context of advanced chronological age, was related to augmented WM aging (i.e. BAG) supports the first hypothesis. In line with this hypothesis, HIV effects on the brain have been shown to occur via similar cellular mechanisms as normal aging^36^, including alterations of the neuroprotective and inflammatory functions of microglia^37^, white matter microvasculature changes^38^, cholinergic deficit^39^ and accumulation of amyloid-beta and tau plaques^40^. Any single or combination of these mechanisms could lead to loss of WM microstructural organization through axonal injury, (in line with AD contributions to SVR model), loss of axonal density (MD contributions) and/or reduced patency of axons (RD contributions).

To the best of our knowledge, this is the first study to demonstrate an HIV-associated accentuated aging process in WM microstructure, using DTI. These findings are generally consistent with the literature, including a recently published study showing an HIV-related accentuated aging process in combined grey matter/white matter volume^9^. Similar to that reported by Cole et al., we did not find significant relationships between WM BAG and Nadir CD4 or HIV duration. Our findings expand upon these previous results by providing data suggesting the mechanism through which this augmented aging process deleteriously affects WM microstructure. Our results also further the literature in that we found a significant relationship between BAG and peak HIV RNA viral load as well as a significant age*peak viral load interactive effect on BAG, indicating that the effects of disease burden on brain integrity are more pronounced with advanced age.

Further, these findings indicate the significant advantages of using BAG to predict HIV-associated white matter aging over other methods. The BAG findings were much stronger than our conventional age-trajectory findings, indicating that the SVR-based brain age approach we used is a sensitive approach to reveal group differences beyond simple differences in mean DTI measures. Additionally, BAG outperformed each individual DTI metric in its ability to discriminate between HIV+ and HIV-participants and demonstrate the effect of HIV infection on advancing brain white matter age. BAG also may be more useful than, or at the very least a meaningful compliment to, hyper/hypo-intense lesion volume and count which has been shown to relate to HIV infection and cognitive performance, but not to HIV clinical variables or HIV-associated aging^41, 42, 43^. Additionally, BAG is a relatively easy metric to understand and thus it circumvents the cumbersome and difficult to interpret multivariate score often used with DTI metrics which are inherently difficult to connect to clinical variables. In contrast, BAG was successfully and clearly connected to both HIV clinical variables (e.g. HIV RNA viral load) and neurocognitive performance.

It is important to consider these findings also in the context of the psychosocial stressors and associated comorbidities associated with living with HIV. For example, the HIV+ sample evidenced greater rates of comorbidities, both medical (which were included in the model herein) and psychiatric (e.g. depression, which was not included in the model). It is unclear in the literature to what extent depression is a secondary reaction to living with HIV or is a neurologic symptom of the predominantly frontal-subcortical clinical profile of the disease. Therefore, it remains unclear whether the augmented aging findings are related directly and solely to the effects of the HIV virus on the brain or if they are also related to secondary effects of these HIV-associated increased comorbidities. This is particularly worthy of follow up investigation given that depressive disorders are the most prevalent mental health disorders associated with HIV^44^ and studies have shown that depression can be associated with neurodegeneration and increased brain age^45^.

There are limitations of the current study worth noting. First, the cross-sectional nature of this study hinders our ability to make inferences about the rates of neuroanatomic changes in HIV. Importantly, this limited our ability to determine whether our findings relate to a more static, vestigial process which adds to or augments the aging process in HIV, or whether in fact these results are related to a dynamic, *accelerated* aging process. This is an important distinction and clinically meaningful question, particularly as the HIV+ population continues to age in the post-HAART era, and must be addressed using a longitudinal model. A recent longitudinal publication^45^ demonstrated that HIV+ participants demonstrated greater predicted brain age than HIV-controls when analyzed at cross-section. However, when followed longitudinally, the HIV+ and HIV-groups evidenced comparable rates of change in neuroimaging markers, suggesting that, when receiving successful treatment, people living with HIV are not at risk for accelerated brain aging over two years. Longer longitudinal studies will help clarify whether or not this pattern remains steady over time.

Next, the SVR-modeling of the DTI data appeared to be less accurate (MAE = 7.39 years) than that using T1-MRI to measure brain volume (MAE = 5.01 years)^9^. This could be due to differences in the neuroimaging methodology used (e.g. size, variability and number of features of training sample set). However, it is also the case that we sought to test a different biological entity (DTI-based WM microstructure), and as such a direct comparison between SVR-derived brain ages may not be appropriate, as Cole et al sought to determine a best-predicted brain age based on grey/white matter volume and we sought to determine the best-predicted age of WM microstructure‥ The fact that data was acquired at multiple sites using different MR scanner could be a factor and a limitation. However, scanner was included as a variable in the model and the data was homogenized using a single, uniform processing pipeline which has become an accepted standard of practice in the field and indeed has been used in similar machine learning papers^9, 10^ where the training data and the disease-specific data were collected at separate sites using different scanners and non-identical scanning parameters. Therefore, this is a limitation of note but one whose impact on the findings was minimized to the best of our abilities. Additionally, although the TBSS method applied should limit the impact of atrophy on our findings, this study did not employ any specific control regarding possible WM lesions. Given that WM lesions have been reported in the brains of HIV+ patients, it is possible that our prediction of WM age could be improved had we included WM lesions from any affected patient in the model. Further, there are some inherent demographic differences (e.g. race/ethnicity) in the training datasets (from England and Norway) and the test dataset (from Los Angeles). While we attempted to control for these by comparing our UCLA HIV+ participants to UCLA HIV-participants who were highly matched on demographic variables and assessing the effect of race/ethnicity on the outcomes, it remains possible that such demographic or even genetic variables could contribute to our findings, though we believe this is less likely for several reasons including the SVR model fit statistics and similar findings from Cole et al.^9^ Further, there are some limits to the generalizability of this study. These include the exclusion of participants with substance use disorders and Axis I diagnoses, the (although non-significant) fact that our sample included fewer HIV+ women, and the fact that the older HIV+ adults are long term survivors from the pre-HAART era and may not be representative of HIV+ adults reaching older age in the near future who were diagnosed in the post-HAART era. Importantly though, it is possible that comorbid substance abuse and/or psychiatric disorders may increase the risk of premature brain aging. Although the current CD4+ and viral load data used in this study were extracted from blood samples collected during the course of this study, nadir CD4+ and highest lifetime viral load were self-reported by participants. Additionally, the DTI-based WM metrics were extracted from the entire white matter skeleton, rather than from individual WM tracts (e.g. uncinate fasciculus). This method limits the spatial resolution of our technique. Future examinations between specific tracts with respect to white matter aging are warranted.

Despite these limitations, the findings of the current study support the hypothesis of HIV-associated augmented brain aging and provide a unique contribution to the existing literature by demonstrating that the mechanism by which this process occurs in WM microstructure appears to be related to HIV-associated neurodegeneration, including axonal injury, loss of axonal density and reduced patency of axons that likely occurs via similar cellular mechanisms as typical aging. Importantly, neuroimage-derived age predictors may indeed be biomarkers of normal and pathologic aging processes. Therefore, this technique may be generalizable to other disease processes which may affect the aging process, including neurodegenerative disorders (e.g. Alzheimer’s disease) and other neuro-medical illnesses. This technique may also be useful in identifying patients at risk for cognitive decline, functional limitations, and early mortality^10^.

## ACKNOWLEDGEMENTS

This work was supported by the University of California, Los Angeles Neurologic, Neurocognitive and Functional Consequences of HIV K23 MH095661 (awarded to A.D.T.) and the Clinical and Translational Research Center Grants UL1RR033176 and UL1TR000124 (awarded to A.D.T.). This work was also supported by the Mount Sinai Institute for NeuroAIDS Disparities NIMH R25 MH080663 (pilot grant awarded to T.P.K.). University of Oslo authors (L.T.W., T.K., N.T.D.) were funded by the Research Council of Norway (213837, 223273, 204966/F20, 229129, 249795/F20), the South-Eastern Norway Regional Health Authority (2013-123, 2014-097, 2015-073), the KG Jebsen Foundation, the European Commission 7th Framework Programme (602450, IMAGEMEND). Taylor Kuhn, Ph.D. is supported by an NIH T32 Training Grant (MH19535). Dr. Singer is supported by U24MH100929. Data collection and sharing for this project was provided by the Cambridge Centre for Ageing and Neuroscience (CamCAN; available at http://www.mrc-cbu.cam.ac.uk/datasets/camcan/)^13, 14^. CamCAN funding was provided by the UK Biotechnology and Biological Sciences Research Council (grant number BB/H008217/1), together with support from the UK Medical Research Council and University of Cambridge, UK. We would like to thank Ariana Anderson, Ph.D., for her contributions during the statistical planning and manuscript preparation stages. The authors report no disclosure.

